# Deep protein representations enable recombinant protein expression prediction

**DOI:** 10.1101/2021.05.13.443426

**Authors:** Hannah-Marie Martiny, Jose Juan Almagro Armenteros, Alexander Rosenberg Johansen, Jesper Salomon, Henrik Nielsen

## Abstract

A crucial process in the production of industrial enzymes is recombinant gene expression, which aims to induce enzyme overexpression of the genes in a host microbe. Current approaches for securing overexpression rely on molecular tools such as adjusting the recombinant expression vector, adjusting cultivation conditions, or performing codon optimizations. However, such strategies are time-consuming, and an alternative strategy would be to select genes for better compatibility with the recombinant host. Several methods for predicting soluble expression are available; however, they are all optimized for the expression host *Escherichia coli* and do not consider the possibility of an expressed protein not being soluble. We show that these tools are not suited for predicting expression potential in the industrially important host *Bacillus subtilis*. Instead, we build a *B. subtilis*-specific machine learning model for expressibility prediction. Given millions of unlabelled proteins and a small labeled dataset, we can successfully train such a predictive model. The unlabeled proteins provide a performance boost relative to using amino acid frequencies of the labeled proteins as input. On average, we obtain a modest performance of 0.64 area-under-the-curve (AUC) and 0.2 Matthews correlation coefficient (MCC). However, we find that this is sufficient for the prioritization of expression candidates for high-throughput studies. Moreover, the predicted class probabilities are correlated with expression levels. A number of features related to protein expression, including base frequencies and solubility, are captured by the model.

## 1. Introduction

Enzymes are the natural catalysts of biochemical processes in every living cell. Understanding the expression of enzymes is a crucial step in engineering them for biotechnological applications (Madigan et al., 2003). Industrial production of enzymes requires recombinant expression in a host microbe under favorable conditions. However, the expression of enzymes is an art form, and large amounts of effort and resources are needed for it to succeed (Habibi et al., 2014).

Multiple factors are known to influence the outcome of recombinant protein production. These include codon usage of the gene (Fu et al., 2020), expression vector and plasmid design (Rosano and Germán, 2019), host strain design and optimizations, growth media, and cultivation conditions, as well as protein recovery method (Zhang et al., 2020). In addition, some proteins can be toxic to the host or aggregate in inclusion bodies (Rosano and Germán, 2019). To ensure a robust expression system, variability in the above factors must be minimized, such as keeping the host strain, expression vector, and growth conditions constant. However, due to the variation in natural proteins, this is not always possible. To handle the variations, multiple growth media and cultivation conditions can be explored, as can optimizations of the gene’s codon usage to better match the codon usage of the recombinant host (Fu et al., 2020). The structure of the mRNA transcript is also known to influence gene expression, which can be optimized by comparing folding energies (Kudla et al., 2009; Cambray et al., 2018). The above factors and variability in the expression system are expected to have a significant impact on the protein expression outcome, and strategies for selecting genes more likely to express are needed.

Instead of using the trial-and-error approach to get enough protein overexpression, tools that can direct the selection of genes with a higher probability of successful overexpression are desirable. Several tools have been developed for the prediction of soluble overexpression in *E. coli*, including PROSO II (Smialowski et al., 2012), PaRSnIP (Rawi et al., 2018), DeepSol (Khurana et al., 2018), SKADE (Raimondi et al., 2020), and SoluProt (Hon et al., 2021). In addition, some tools exist for the more specific prediction of solubility, which is an important element in soluble protein expression. These include Protein-Sol (Hebditch et al., 2017) and SoDoPe (Bhandari et al., 2020). The mentioned tools use the primary structure as input and calculate various sequence-based features (e.g., hydrophobicity, charge, k-mer frequencies, disorder), and they use various machine learning techniques: support vector machines (Agostini et al., 2014), gradient boosting machines (Rawi et al., 2018; Hon et al., 2021), neural networks (Khurana et al., 2018; Raimondi et al., 2020), or other statistical methods (Smialowski et al., 2012; Hebditch et al., 2017; Bhandari et al., 2020). However, predicting only soluble proteins can leave out many proteins, ignoring the possibility of a protein being expressed and insoluble (Mehlin et al., 2006). Furthermore, all of these tools have been developed with the host *E. coli* in mind, and it is an open question whether their results can be generalized to other production organisms.

Data-driven tools, especially machine learning (Bishop, 2006), require significant amounts of data. As the cost of sequencing continues to decrease and the number of publicly available data increases (The UniProt Consortium, 2018), machine learning models are becoming better suited to predict protein characteristics and functions (Elnaggar et al., 2020). This, in combination with progress in computational power and access to machine learning frameworks (Abadi et al., 2015; Pedregosa et al., 2012), have led to new data-driven tools for protein modeling tasks (Almagro Armenteros et al., 2017; Bileschi et al., 2019; Rives et al., 2021; Strodthoff et al., 2019; Alley et al., 2019).

However, the majority of available data is unlabeled or labeled by existing prediction methods only and cannot easily be used for learning protein properties with supervised machine learning. In order to utilize the vast amounts of unlabelled data, a method known as UniRep (Alley et al., 2019) has been developed to convert biological sequences into statistical representations. The protein embeddings are able to incorporate structural, evolutionary, and biophysical features despite having no prior information on them. UniRep uses language modeling (Jurafsky and Martin, 2019) to build the representation by predicting which amino acid comes next given the prior sequence.

This study examines a dataset of proteins that have been experimentally validated for expression in the gram-positive bacterium *B. subtilis*, an important production host in the biotechnological industry. This dataset presents a prime opportunity to build a model that can predict the probability of a gene being expressed in *B. subtilis*, based only on the amino acid sequence of the protein. In the recombinant expression system, most molecular parameters such as recombinant host and expression vector were kept constant; a fixed set of growth media and cultivation conditions were explored, and codon optimizations were not performed, using only the codons of the natural gene.

We investigate several modeling approaches on how to solve this prediction task and find that using features generated by UniRep significantly improves performance relative to using amino acid composition. To our knowledge, this is the first time unsupervised learning has been successfully applied to the prediction of recombinant expression.

Furthermore, we show that specific UniRep features correlate with biological features important for protein expression in *B. subtilis*. We demonstrate that universal sequence representations are better suited to capture features important for predicting recombinant gene expression than building a model that is solely trained on host-specific data. The study is summarized in Figure 1.

**Figure 1:**
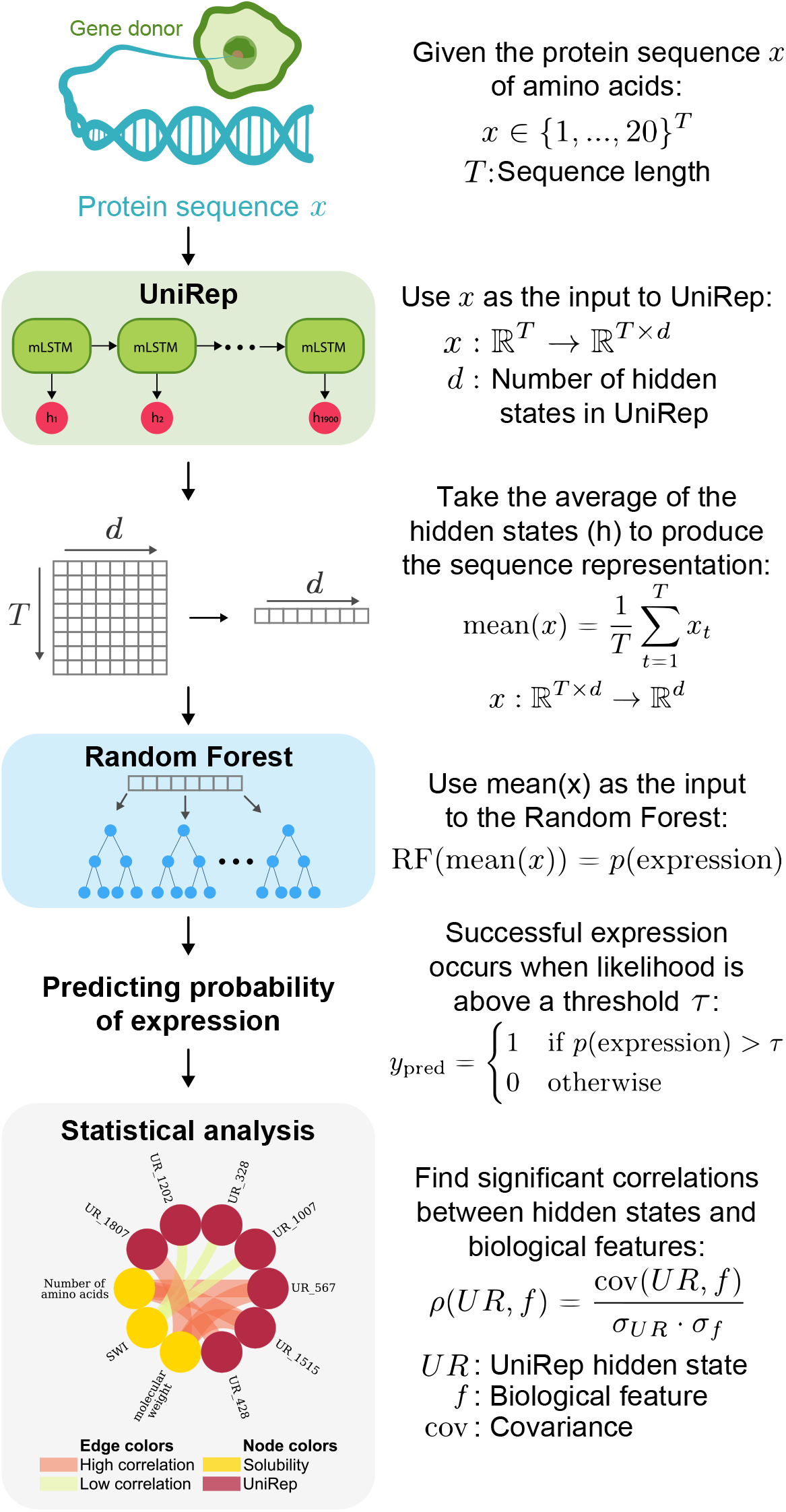
Summary of our work. We have designed a system in which a protein sequence from an organism (gene donor) is converted into a numerical vector by the hidden states of UniRep (Alley et al., 2019). The vector is then used as an input to a classifier, i.e., a random forest, that predicts whether the protein can be recombinantly expressed. Finally, we show that specific hidden states (units) correlate to biological features in a Circos plot (Krzywinski et al., 2009) with nodes being units (UR) and features connected by edges colored and sized by the absolute correlation.

## 2. Materials and methods

We define the problem of predicting recombinant gene expression in *B. subtilis* as a binary classification (success or failure) and evaluate different classifiers by their performance on a held-out test set. We test a range of machine learning classifiers using either amino acid frequencies or the internal states of UniRep as input, and we find that the latter input type significantly improves the performance.

### 2.1. Bacterial expression dataset

The dataset consists of 4487 genes, which have been collected and experimentally tested for expression in *B. subtilis* by Novozymes A/S.

Various methods have been developed to characterize recombinant expression, where the PCR-based cloning approach has been used to verify the proteins in this dataset. Whole coding regions of the bacterial genes were amplified by PCR from genomic DNA and cloned into an expression vector (Widner et al., 2000). The PCR fragment and vector were digested with restriction enzymes. Vector and fragment were ligated, and the recombinant plasmids were used to transform *E. coli* yielding several recombinants per gene. A plasmid containing a confirmed gene sequence was transformed into *B. subtilis* and subsequently one recombinant *B. subtilis* clone containing the integrated expression construct was grown in a liquid culture at temperatures ranging from 20 °C-37 °C for 2-5 days. The cultures were harvested, supernatant extracted and analyzed by SDS-PAGE electrophoresis. The proteins were visualized by staining with Cromassie Blue G-250, and estimation of molecular weights and yields of the proteins were made against molecular (stained) standards (250, 150, 100, 75, 50, 37, 25, 20, 15, 10 kDa). If no visible band of the size of the protein was detected on the SDS-PAGE, a gene was determined to be not expressed. Different dyes and contrast levels can affect the readout of SDS-gels, but typically concentrations of down to 50mg/l will produce visible bands. The sizes of the bands were also used to estimate expression levels for section 3.2.

We use homology partitioning to partition the data into training, validation, and test sets to ensure better generalization of our held-out validation and test set. PSI-CD-HIT (Li and Godzik, 2006) is used to cluster sequences with 30% identity. Based on the homology, the clusters were used to divide the data into 70% training, 10% validation, and 20% test set partitions while maintaining the ratio between successful and unsuccessful expression in each set.

### 2.2. Modeling overview

We test a variety of machine-learning-based tools for their ability to predict recombinant gene expression. We compare the prediction performance of support vector machines (SVM), logistic regression (LR), random forest (RF), and artificial neural network (ANN)(Hastie et al., 2016) using either amino acid frequencies or a pretrained language model (UniRep (Alley et al., 2019)) to embed our proteins before training a model that predicts gene expression. Parameter optimization is done for all classifiers, and each training round was repeated with 10 different random seeds to obtain balanced performance measurements.

#### 2.2.1. Modeling details

Given either the amino acid frequencies or the UniRep embeddings as input, we train an SVM, LR, RF, and ANN to predict recombinant expression. Given the validation set, we perform a hyperparameter optimization as follows; the hyperparameters optimized for the SVM are the scaling term for the regularization used in stochastic gradient descent and the tolerance value in the stopping criterion. The SVM uses a linear kernel and the modified Huber loss function. For LR, we optimized the tolerance value as well as the inverse regularization term value. For the RF, the following hyperparameters are optimized: the number of trees, the number of features for the best split, the minimum number of samples required to make a split, and whether to use bootstrap (Hastie et al., 2016). We optimize the number of hidden layers in the ANN and the number of hidden units in each of the hidden layers. The learning rate and the L2 penalty term are also optimized. Training of the neural network is done with Adam optimization (Kingma and Ba, 2014) and early stopping regularization is used. If a hyperparameter is not listed as being optimized, we use the default values in their implementation in scikit-learn version 0.20.2 (Pedregosa et al., 2012).

#### 2.2.2. UniRep details

UniRep (Alley et al., 2019) takes a protein sequence as input and extracts 1900 continuous features. UniRep is based on a deep learning (LeCun et al., 2015) method known as the recurrent neural network (RNN) (Hochreiter and Schmidhuber, 1997). The RNN is trained on the UniRef50 database containing more than 24 million protein sequences (Suzek et al., 2015). The 1900 units are the averages of the RNN hidden states across the sequence. Our protein sequences are represented with UniRep and then used as inputs to each of the classifiers.

### 2.3. Evaluation

To measure relative performance, corrected for class imbalances, we calculate the area under the receiver operating characteristic (ROC) curve, known as the AUC, and the Matthews correlation coefficient (MCC) (Matthews, 1975).

The ROC curve visualizes a trade-off between the true-positive rate (TPR, sensitivity) and the false-positive rate (FPR, 1-specificity) when increasing the probability threshold for classification, *τ*. The area under the ROC curve (AUC) is used as a summary statistic of the global accuracy of the predictor. A guideline for interpreting the AUC by Swets (1988) indicates that an AUC of 0.5 is similar to random selection (our baseline), and the closer AUC is to 1, the more accurate the predictor is. The best model among the evaluated architectures was chosen based on the AUC.

The MCC is a performance metric that takes into account dataset imbalance (positive samples are overly represented). A random model will achieve a performance of 0.0, and an oracle model would get 1.0.

The behavior of the classifier is highly dependent on the cut-off value, *τ*, since higher values of *τ* will decrease the sensitivity (Se) and increase the specificity (Sp) and vice versa (Greiner et al., 2001). We use the Youden Index *J* = max_*τ*_ {Se(*τ*) + Sp(*τ*) − 1} (Youden, 1950) to select the value of *τ* based on the validation ROC curve, where *J* is set to put equal weight on the sensitivity and specificity of the model (Fluss et al., 2005; Greiner et al., 2001). Classifying a gene to be expressed occurs when the likelihood is above the threshold (*P* (gene) > *τ*), meaning that we expect confident answers are more likely to be correct (Johansen and Socher, 2017).

Finally, we compare our results on the validation and test set with predictors published in earlier studies. We evaluate the following predictors of solubility or soluble expression in *E. coli* on our *B. subtilis* sequences: Protein-Sol (Hebditch et al., 2017), SKADE (Raimondi et al., 2020) and SoDoPe (Bhandari et al., 2020).

### 2.4. Sequence-based feature generation

Features related to protein solubility were generated with Protein-Sol (Hebditch et al., 2017) and SoDoPe (Bhandari et al., 2020). The selected Protein-Sol features were seven amino acid composites (K-R, D-E, K+R, D+E, K+R-D-E, K+R+D+N, and F+W+Y) and eight protein predicted features (protein length, isoelectric point (pI), hydropathy, absolute charge at pH 7, fold propensity, disorder, sequence entropy, and beta-strand propensity). From SoDoPe, we added the predicted solubility score (SWI) for a protein.

Predicted secondary structures of the proteins were made by Porter 4.0 (Mirabello and Pollastri, 2013), which classifies a protein sequence into three classes: Helix, Strand, or Coil. We converted the counts into percentages of helices, strands, and coils for a given sequence.

Information about enzyme type and origin was provided by Novozymes A/S and were added to the set of sequence-based features.

### 2.5. Correlations between UniRep units and biological features

In order to understand which features UniRep captures in its embedding of our protein sequences, we calculated Pearson’s correlation coefficient between sequence-based features and the vectors with UniRep represented sequences and the p-value for testing non-correlations. We report only statistically significant correlations (p-value ≤ 0.05 with Bonferroni correction) for the validation and test data (Hastie et al., 2016). Correlations are done for predicted secondary structures, solubility properties, amino acid frequencies, enzyme, and taxonomic labels.

Correlations between UniRep values and the sequence features are visualized in a heatmap or in Circos plots (Krzywinski et al., 2009), where each node is either a UniRep or a biological feature, and an edge shows the strength of the correlation between two nodes.

### 2.6. Code and data availability

The code for getting UniRep representations for amino acid sequences, training the Random Forest, and correlating features are available at https://github.com/hmmartiny/Predicting-Gene-Expression. Unfortunately, due to corporate confidentiality, we are unable to publish the experimental data but encourage others to try the code on their own data.

## 3. Results

Table 1 contains the results of our comparison of various approaches to predicting recombinant gene expression in *B. subtilis* with the two types of protein inputs. We benchmark the following modeling architectures on their performance of the held-out test set: SVM, LR, RF, and ANN. Furthermore, we evaluate the effect of using amino acid frequencies as input as compared to using UniRep formatted sequences. Performance is measured by the area under the ROC curve (AUC) and Matthew’s correlation coefficient (MCC).

**Table 1:** Validation and test set performances by the various model architectures. AA: amino acid. UniRep sequence: protein sequence represented with UniRep. Model hyper-parameters are in Table A.3 and Table A.4.

| Input | Score | SVM | LR | RF | ANN |
| --- | --- | --- | --- | --- | --- |
| AA frequency | Valid AUC | $0.55 \pm 0.02$ | $0.57 \pm 0.00$ | $0.57 \pm 0.01$ | $0.58 \pm 0.01$ |
| | Test AUC | $0.58 \pm 0.02$ | $0.60 \pm 0.00$ | $0.59 \pm 0.01$ | $0.59 \pm 0.01$ |
| | Test MCC | $0.13 \pm 0.05$ | $0.15 \pm 0.00$ | $0.10 \pm 0.02$ | $0.15 \pm 0.02$ |
| UniRep sequence | Valid AUC | $0.59 \pm 0.02$ | $0.60 \pm 0.00$ | $0.66 \pm 0.01$ | $0.62 \pm 0.01$ |
| | Test AUC | $0.62 \pm 0.02$ | $0.64 \pm 0.00$ | $0.64 \pm 0.01$ | $0.64 \pm 0.01$ |
| | Test MCC | $0.15 \pm 0.04$ | $0.20 \pm 0.00$ | $0.20 \pm 0.02$ | $0.20 \pm 0.03$ |

Our results show that using UniRep to format the sequences boosts performances with the ANN scoring test 0.64 ± 0.01 AUC and 0.20 ± 0.03 MCC for *τ* = 0.60 and the RF achieving 0.64 ± 0.01 AUC and 0.20 ± 0.02 MCC for *τ* = 0.61. Interestingly, we find that using either input is better than random guessing but that UniRep formatted sequences gave the highest performance. Comparing our models with existing frameworks (Protein-Sol (Hebditch et al., 2017), SKADE (Raimondi et al., 2020) and SoDoPe (Bhandari et al., 2020)) show that models built on data coming from one type of host organism (i.e. *E. coli*) are not comparable to the universal embeddings learned by UniRep (Table 2). The performance gained by using UniRep based models indicates that unsupervised feature extraction can be useful for predicting recombinant gene expression in different production organisms.

**Table 2:**
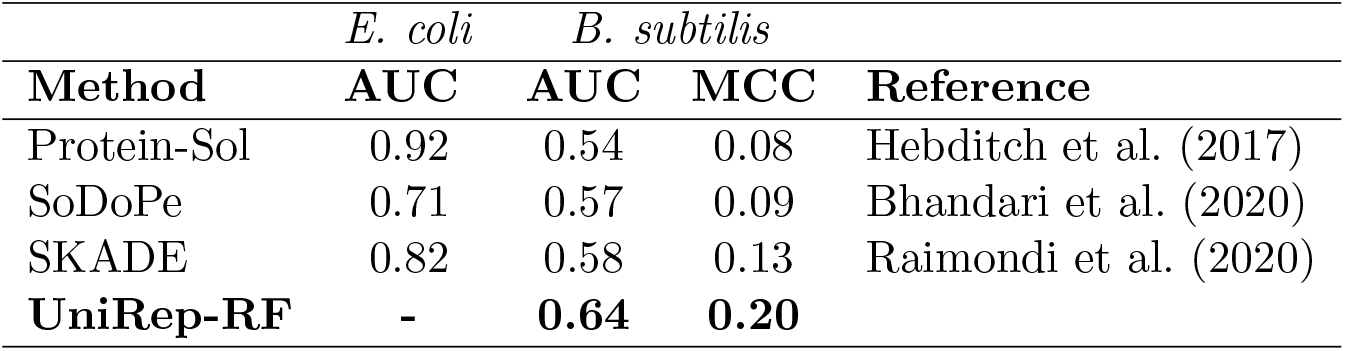
Comparison of selected solubility predictors performance on *E. coli* test sets and our *B. subtilis* test set. The reported *E. coli* AUC scores are those reported on the test data used in the papers.

In Figure 2, we analyze what happens when only considering samples above a certain probability threshold in order to maximize specificity. Our results suggest that the model has increased precision on proteins with high confidence. E.g., thresholding the probability of expression to *τ* = 0.8 gives a precision of more than 80%. However, this reduces the amount of available samples to less than 10% of the test samples.

**Figure 2:**
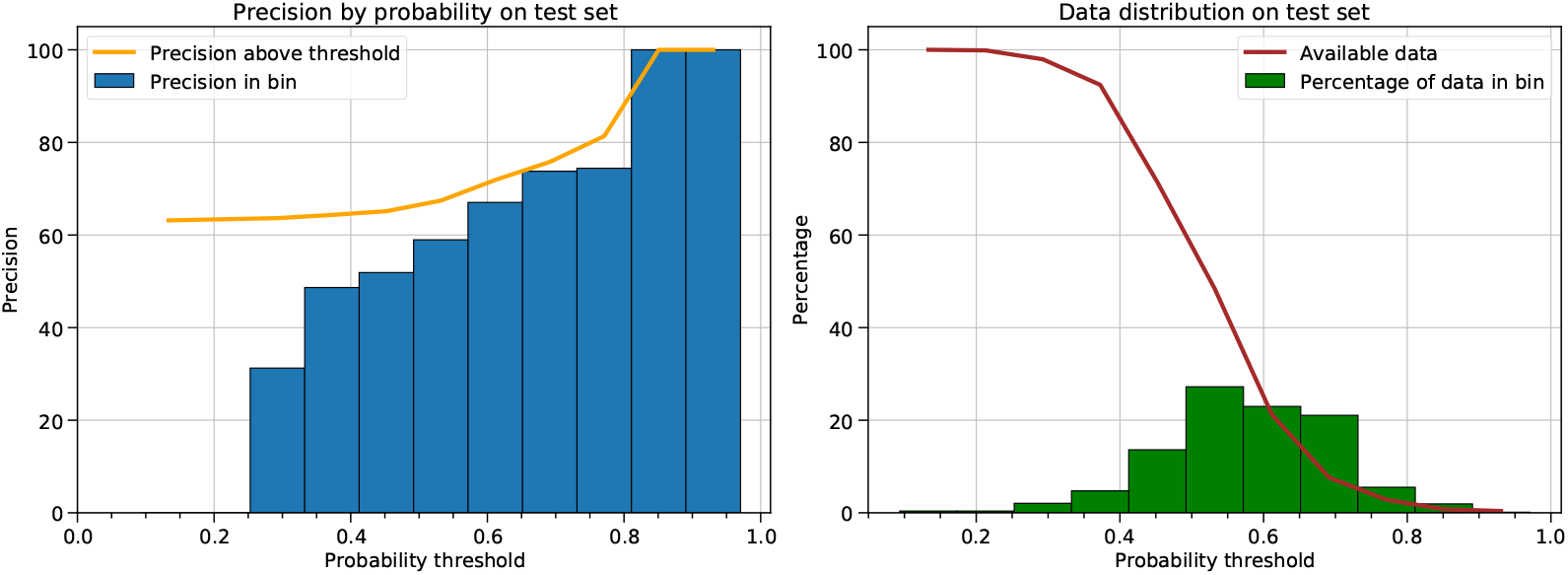
Precision and data distribution by UniRep-RF probability thresholds on the test set. Left: Blue bars correspond to precision by each bin, and the orange curve is the precision of all samples above the threshold. The precision is calculated by saying that all the samples in a bin are positive. Right: Green bars show the percentage of data in each bucket, and the brown curve corresponds to the amount of available data above a threshold.

### 3.1. Specific UniRep units correlate to protein features

In an attempt to understand why using UniRep formatted sequences raised test performances, we examine the correlation between each element of the 1900-unit vector produced by UniRep for each protein sequence and various protein features (Figure B.5). Pearson’s correlation coefficient is used, and we selected only statistically significant values (p-value < 0.05 with Bonferroni correction).

Only 25 out of the 1900 units are repeatedly selected as being part of the 10 most important features in the differently seeded UniRep-RFs, and their correlation to protein features in the test set are shown in Figure 3. Feature importance is measured as the Gini impurity (Hastie et al., 2016), meaning the decrease in node impurity for each feature. It can be seen that the 25 units can be clustered into two large groups, with one being a small-sized cluster that contains 7 units that correlate to many of the protein features. Especially unit 932 seems to capture base frequencies (amino acid and nucleotide) in the sequences as well as protein properties (e.g., secondary structure or solubility), although many of the 25 units have varying levels of correlations to the latter. See Figure B.5 for which UniRep units correlate to which protein features.

**Figure 3:**
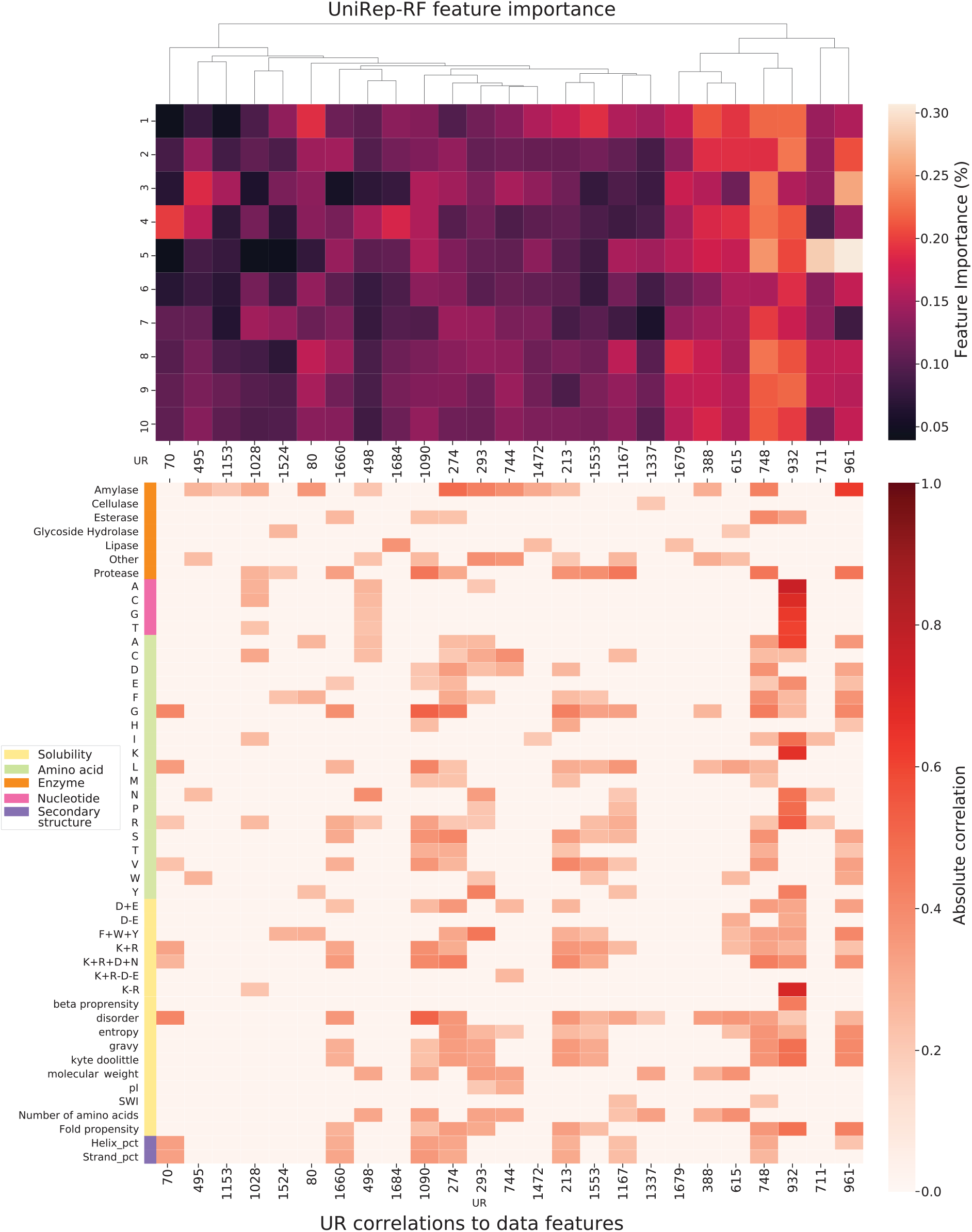
The most important UniRep (UR) units for the RFs and what they correlate to. Top: Heatmap visualization of feature importance (%) of all selected UR that are in the top 10 most important (%) in the differently seeded RFs (1–10). Bottom: Heatmap showing the correlations between a UR and a data feature. A zero correlation indicates that the correlation is not statistically significant (p-value > 0.05).

### 3.2. Prediction scores correlate with estimated expression levels

Following the development of the model, 108 additional genes were expressed. Expression yields were categorized into Low (estimated 0.05-0.15 g/l), Medium (0.2-0.5 g/l), and High yields (>0.5 g/l) by relative band sizes. Figure 4 shows a box plot of the estimated expression yields vs. the expressibility likelihood score per yield category.

**Figure 4:**
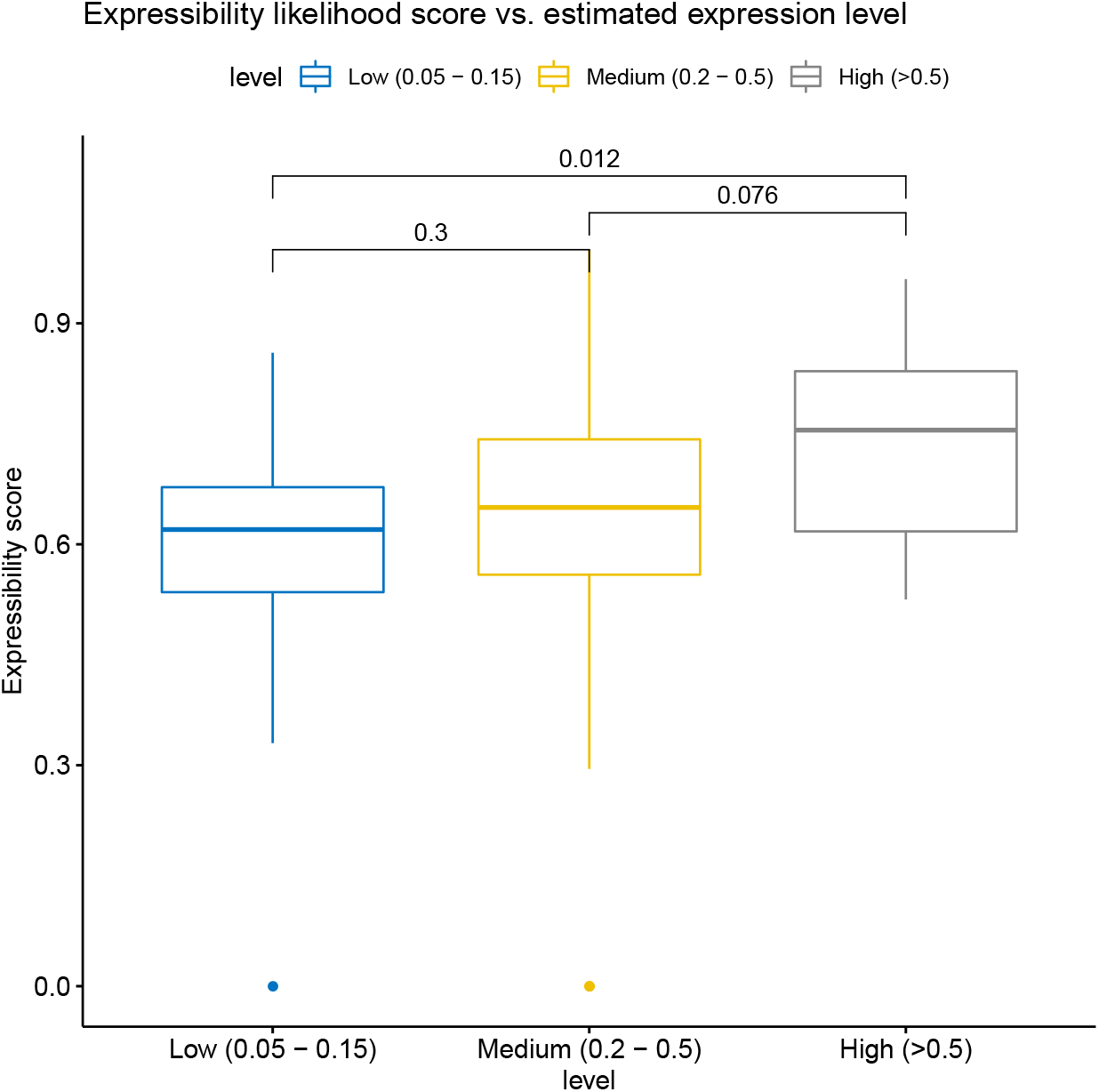
Box plot of categorized estimated expression levels and expressibility likelihood scores of 108 expressed genes. P-values of t-tests between groups are shown at the top.

## 4. Discussion

Testing expressibility of a protein in a production host typically entails several weeks of lab work and only one outcome (success or failure). The outcome may depend on several factors extrinsic to the amino acid sequence, such as experimental conditions and codon usage. Despite this inherent noise in the data, we find that UniRep features, combined with a non-linear classifier, can extract generalizable information from the training set and deliver predictions that are better than the composition-based baseline.

Our approach is well suited for screening a large number of proteins in high-throughput studies since the model makes it possible to reduce the number of proteins to test experimentally by thresholding the likelihood of expression. Although the performance of the UniRep-RF model is not the most impressive, focusing on high likelihood proteins would result, with statistical significance, in higher expression rates.

Neither unsupervised learning nor Random Forests models are novel methods, but the use of both to predict the expression of proteins is new. There are many reasons why a protein can or can not be expressed in a host. Still, we show that a small set of UniRep units correlate to biochemical and taxonomic features, confirming the use of unsupervised learning techniques for protein engineering tasks.

The small subset of UniRep units that correlate to selected protein features could be used to inform which features to optimize for improved expression. To achieve this, more work is needed to verify which units capture what feature and include the units that were not part of the 25 important ones for the classifier. This could involve correlating a large set of enzyme sequences, regardless of whether they have been tested for recombinant expression in *B. subtilis*. Since there are many reasons as to why a protein is or is not expressed, several other properties, such as mRNA folding, could be correlated as well.

UniRep learned protein embeddings by training on amino acid sequences from all aspects of life, so our approach could be expanded to include other expression systems than *B. subtilis*. UniRep is not the only published model that can extract information-rich sequence representations, so comparing the relative performances of other pretrained models, such as Strodthoff et al. (2019), Rives et al. (2021) or Brandes et al. (2021), might reveal other important features.

## Appendix A. Parameters for classifiers

**Table A.3:**
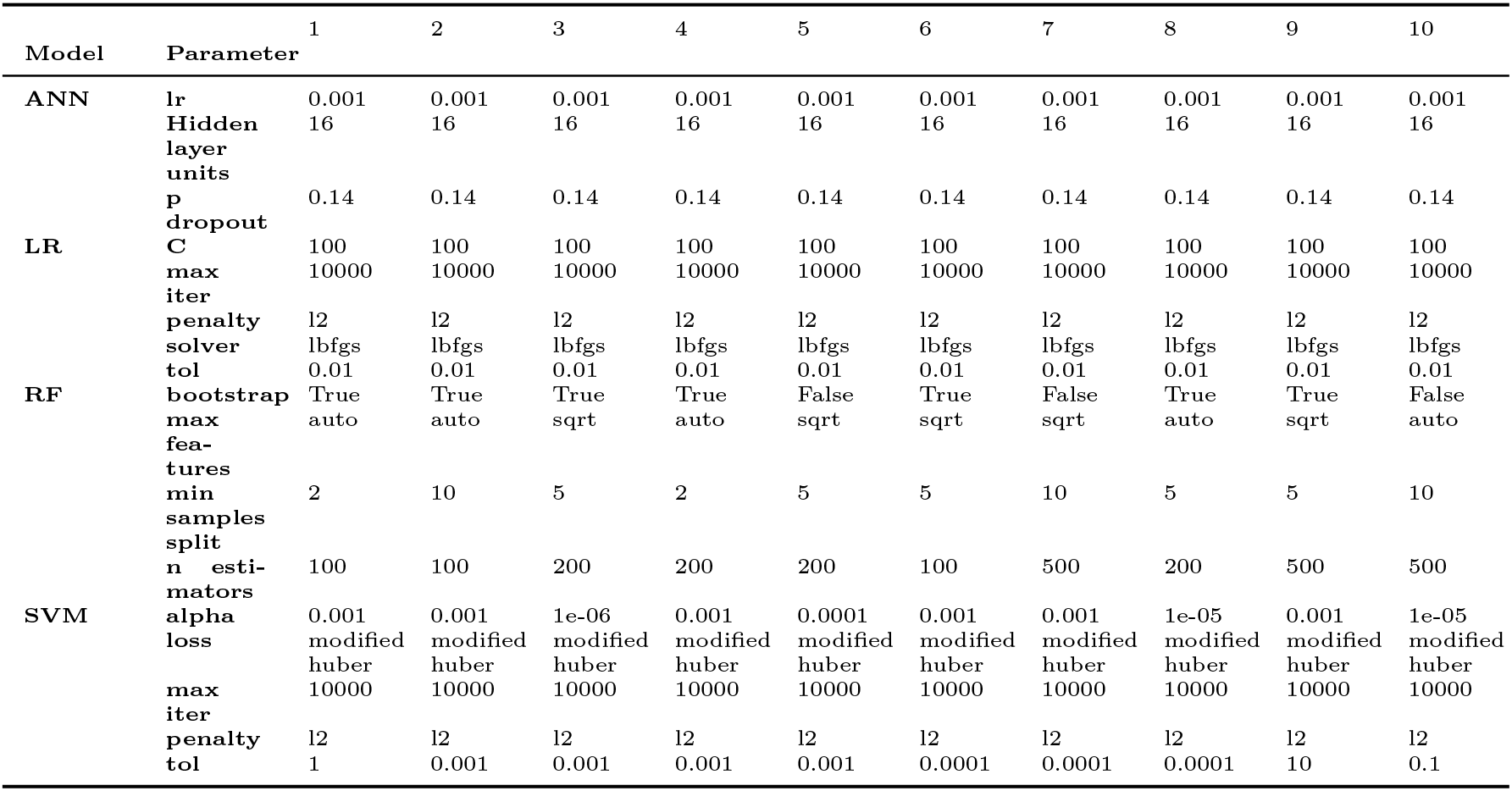
Parameters for AA Frequency classifiers

**Table A.4:**
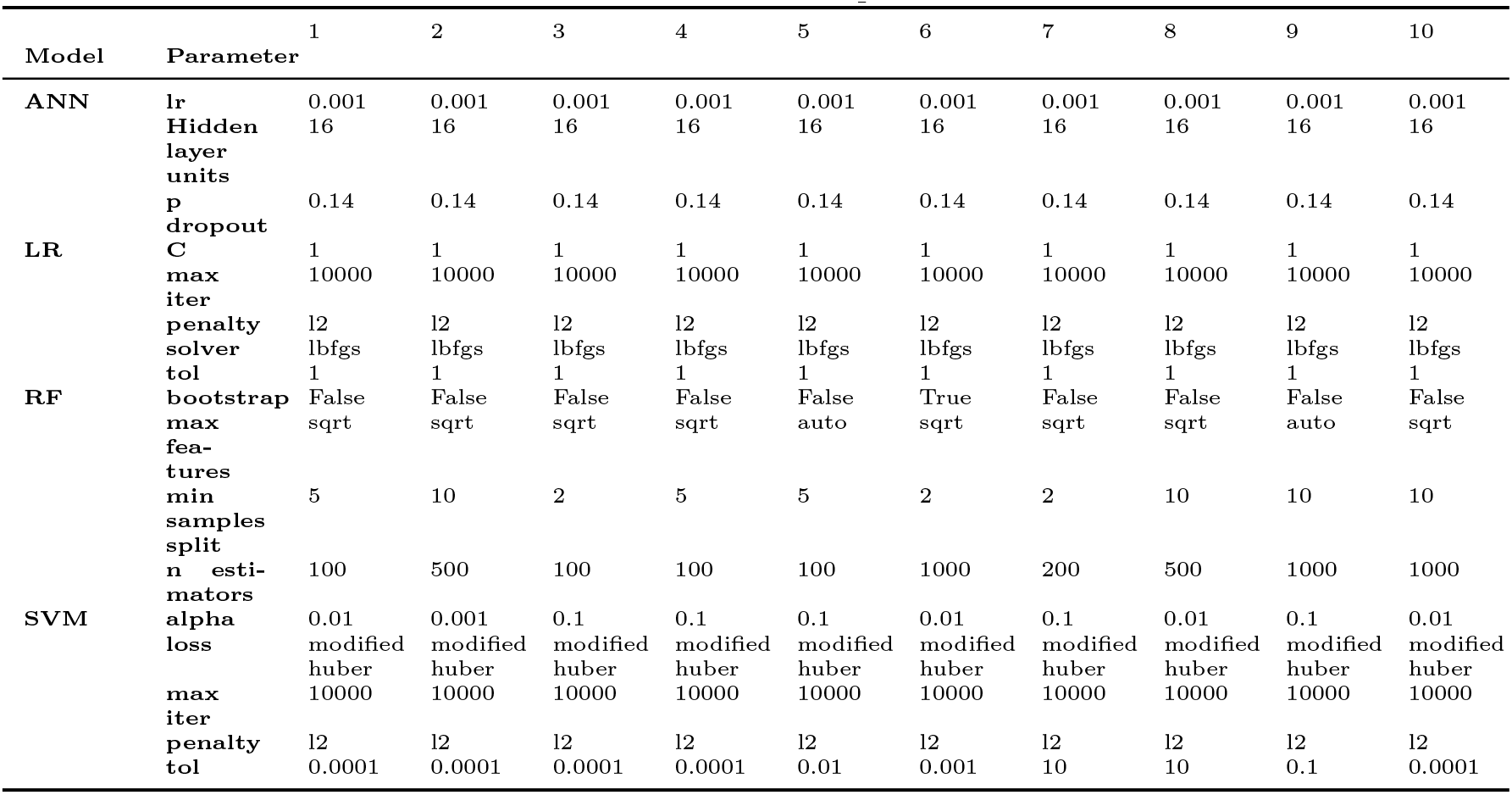
Parameters for UniRep classifiers

## Appendix B. Correlations

**Figure B.5:**
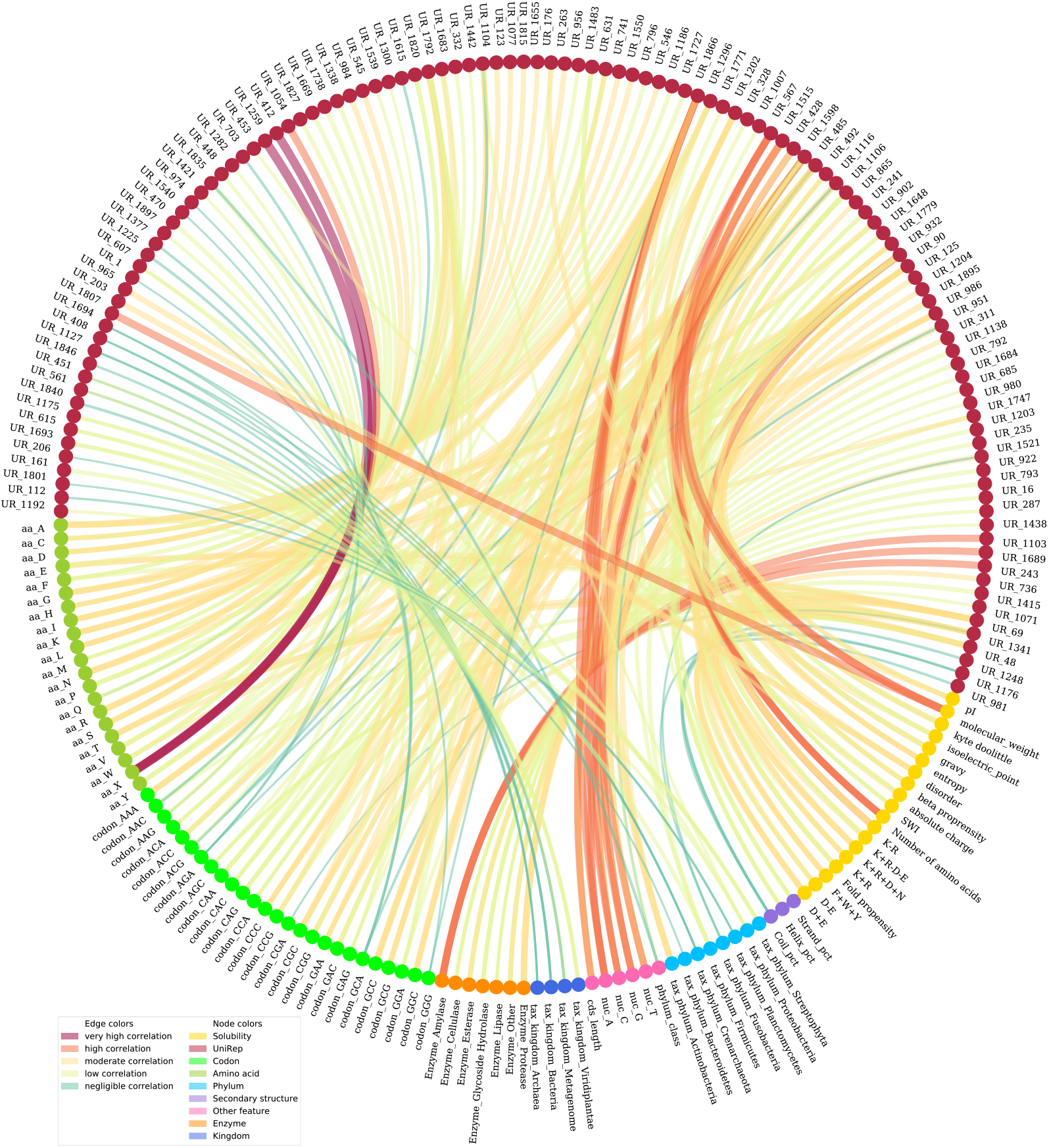
Circos plot showing the three highest correlations between each pair of all sequence based features and UniRep units (UR). A node is either a UR or a feature connected with an edge showing the strength of the absolute correlation. Only correlations that were statistically significant are shown.

## References

Abadi, M., Agarwal, A., Barham, P., Brevdo, E., Chen, Z., Citro, C., Corrado, G. S., Davis, A., Dean, J., Devin, M., Ghemawat, S., Goodfellow, I., Harp, A., Irving, G., Isard, M., Jia, Y., Jozefowicz, R., Kaiser, L., Kudlur, M., Levenberg, J., Mané, D., Monga, R., Moore, S., Murray, D., Olah, C., Schuster, M., Shlens, J., Steiner, B., Sutskever, I., Talwar, K., Tucker, P., Vanhoucke, V., Vasudevan, V., Viégas, F., Vinyals, O., Warden, P., Wattenberg, M., Wicke, M., Yu, Y., Zheng, X., 2015. TensorFlow: Large-scale machine learning on heterogeneous systems. Software available from tensorflow.org. URL https://www.tensorflow.org/

Agostini, F., Cirillo, D., Livi, C. M., Delli Ponti, R., Tartaglia, G. G., 2014. cc SOL omics: A webserver for solubility prediction of endogenous and heterologous expression in Escherichia coli. Bioinformatics 30 (20), 2975–2977.

Alley, E. C., Khimulya, G., Biswas, S., AlQuraishi, M., Church, G. M., 2019. Unified rational protein engineering with sequence-based deep representation learning. Nature Methods 16, 1315–1322.

Almagro Armenteros, J. J., Sønderby, C. K., Sønderby, S. K., Nielsen, H., Winther, O., 2017. DeepLoc: prediction of protein subcellular localization using deep learning. Bioinformatics 33 (21), 3387–3395.

Bhandari, B. K., Gardner, P. P., Lim, C. S., 2020. Solubility-weighted index: fast and accurate prediction of protein solubility. Bioinformatics 36 (18), 4691–4698.

Bileschi, M. L., Belanger, D., Bryant, D., Sanderson, T., 2019. Using Deep Learning to Annotate the Protein Universe. bioRxiv, 626507.

Bishop, C. M., 2006. Pattern Recognition and Machine Learning. Springer.

Brandes, N., Ofer, D., Peleg, Y., Rappoport, N., Linial, M., 2021. Proteinbert: A universal deep-learning model of protein sequence and function. bioRxiv, 2021.05.24.445464.

Cambray, G., Guimaraes, J. C., Arkin, A. P., 2018. Evaluation of 244,000 synthetic sequences reveals design principles to optimize translation in Escherichia coli. Nature Biotechnology 36 (10), 1005–1015.

Elnaggar, A., Heinzinger, M., Dallago, C., Rihawi, G., Wang, Y., Jones, L., Gibbs, T., Feher, T., Angerer, C., Bhowmik, D., Rost, B., 2020. ProtTrans: Towards cracking the language of life’s code through self-supervised deep learning and high performance computing. bioRxiv, 2020.07.12.199554.

Fluss, R., Faraggi, D., Reiser, B., 2005. Estimation of the Youden Index and its associated cutoff point. Biometrical Journal 47 (4), 458–472.

Fu, H., Liang, Y., Zhong, X., Pan, Z., Huang, L., Zhang, H., Xu, Y., Zhou, W., Liu, Z., 2020. Codon optimization with deep learning to enhance protein expression. Scientific Reports 10 (1), 17617.

Greiner, M., Pfeiffer, D., Smith, R. D., 2001. Principles and practical application of the receiver-operating characteristic analysis for diagnostic tests. Preventive Veterinary Medicine 45 (2000).

Habibi, N., Mohd Hashim, S. Z., Norouzi, A., Samian, M. R., 2014. A review of machine learning methods to predict the solubility of overexpressed recombinant proteins in Escherichia coli. BMC Bioinformatics 15 (1).

Hastie, T., Tibshirani, R., Friedman, J. H. J. H., 2016. The elements of statistical learning: data mining, inference, and prediction. New York, NY: Springer.

Hebditch, M., Carballo-Amador, M. A., Charonis, S., Curtis, R., Warwicker, J., 2017. Protein–sol: a web tool for predicting protein solubility from sequence. Bioinformatics 33 (19), 3098–3100.

Hochreiter, S., Schmidhuber, J., 1997. Long Short-Term Memory. Neural Computation 9 (8), 1735–1780.

Hon, J., Marusiak, M., Martinek, T., Kunka, A., Zendulka, J., Bednar, D., Damborsky, J., 2021. Soluprot: Prediction of soluble protein expression in Escherichia coli. Bioinformatics 37 (1), 23–28.

Johansen, A., Socher, R., Aug. 2017. Learning when to skim and when to read. In: Proceedings of the 2nd Workshop on Representation Learning for NLP. Association for Computational Linguistics, Vancouver, Canada, pp. 257–264. URL https://www.aclweb.org/anthology/W17-2631

Jurafsky, D., Martin, J., 2019. Speech and Language Processing (3rd Edition). Prentice Hall.

Khurana, S., Rawi, R., Kunji, K., Chuang, G. Y., Bensmail, H., Mall, R., 2018. DeepSol: A deep learning framework for sequence-based protein solubility prediction. Bioinformatics 34 (15), 2605–2613.

Kingma, D. P., Ba, J., 2014. Adam: A Method for Stochastic Optimization. arXiv preprint, 1412.6980.

Krzywinski, M. I., Schein, J. E., Birol, I., Connors, J., Gascoyne, R., Horsman, D., Jones, S. J., Marra, M. A., 2009. Circos: An information aesthetic for comparative genomics. Genome Research 19 (9), 1639–1645.

Kudla, G., Murray, A. W., Tollervey, D., Plotkin, J. B., 2009. Coding-sequence determinants of gene expression in Escherichia coli. Science 324 (5924), 255–258.

LeCun, Y., Bengio, Y., Hinton, G. E., 2015. Deep learning. Nature 521 (7553), 436–444. URL https://doi.org/10.1038/nature14539

Li, W., Godzik, A., 2006. Cd-hit: A fast program for clustering and comparing large sets of protein or nucleotide sequences. Bioinformatics 22 (13), 1658–1659.

Madigan, M. T., Martinko, J. M., Parker, J., 2003. Brock Biology of Microorganisms, 14th Edition. Pearson.

Matthews, B., 1975. Comparison of the predicted and observed secondary structure of T4 phage lysozyme. Biochimica et Biophysica Acta (BBA) - Protein Structure 405 (2), 442–451.

Mehlin, C., Boni, E., Buckner, F. S., Engel, L., Feist, T., Gelb, M. H., Haji, L., Kim, D., Liu, C., Mueller, N., et al., 2006. Heterologous expression of proteins from Plasmodium falciparum: results from 1000 genes. Molecular and Biochemical Parasitology 148 (2), 144–160.

Mirabello, C., Pollastri, G., jun 2013. Porter, PaleAle 4.0: high-accuracy prediction of protein secondary structure and relative solvent accessibility. Bioinformatics 29 (16), 2056–2058.

Pedregosa, F., Varoquaux, G., Gramfort, A., Michel, V., Thirion, B., Grisel, O., Blondel, M., Müller, A., Nothman, J., Louppe, G., Prettenhofer, P., Weiss, R., Dubourg, V., Vanderplas, J., Passos, A., Cournapeau, D., Brucher, M., Perrot, M., Duchesnay, É., 2012. Scikit-learn: Machine learning in python. Journal of Machine Learning Research 12, 2825–2830.

Raimondi, D., Orlando, G., Fariselli, P., Moreau, Y., 2020. Insight into the protein solubility driving forces with neural attention. PLoS Computational Biology 16 (4), e1007722.

Rawi, R., Mall, R., Kunji, K., Shen, C.-H., Kwong, P. D., Chuang, G.-Y., 2018. PaRSnIP: sequence-based protein solubility prediction using gradient boosting machine. Bioinformatics 34 (7), 1092–1098.

Rives, A., Meier, J., Sercu, T., Goyal, S., Lin, Z., Liu, J., Guo, D., Ott, M., Zitnick, C. L., Ma, J., et al., 2021. Biological structure and function emerge from scaling unsupervised learning to 250 million protein sequences. Proceedings of the National Academy of Sciences 118 (15), e2016239118.

Rosano, Germán, 2019. New tools for recombinant protein production in Escherichia coli: A 5-year update. Protein Science 28 (8), 1412–1422.

Smialowski, P., Doose, G., Torkler, P., Kaufmann, S., Frishman, D., 2012. Proso ii–a new method for protein solubility prediction. The FEBS journal 279 (12), 2192–2200.

Strodthoff, N., Wagner, P., Wenzel, M., Samek, W., 2019. Universal Deep Sequence Models for Protein Classification. bioRxiv, 704874.

Suzek, B. E., Wang, Y., Huang, H., McGarvey, P. B., Wu, C. H., 2015. UniRef clusters: A comprehensive and scalable alternative for improving sequence similarity searches. Bioinformatics 31 (6), 926–932.

Swets, J. A., 1988. Measuring the Accuracy of Diagnostic Systems. Science 240 (4857), 1285–1293.

The UniProt Consortium, 2018. UniProt: a worldwide hub of protein knowledge. Nucleic Acids Research 47 (D1), D506–D515.

Widner, B., Thomas, M., Sternberg, D., Lammon, D., Behr, R., Sloma, A., 2000. Development of marker-free strains of Bacillus subtilis capable of secreting high levels of industrial enzymes. Journal of Industrial Microbiology and Biotechnology 25 (4), 204–212.

Youden, W. J., 1950. Index for rating diagnostic tests. Cancer 3 (1), 32–35.

Zhang, K., Su, L., Wu, J., 2020. Recent advances in recombinant protein production by Bacillus subtilis. Annual Review of Food Science and Technology 11 (1), 295–318.

